# Horizontally acquired *CSP* genes contribute to wheat adaptation and improvement

**DOI:** 10.1101/2024.06.04.597356

**Authors:** Kai Wang, Guanghui Guo, Shenglong Bai, Jianchao Ma, Zhen Zhang, Zeyu Xing, Wei Wang, Hao Li, Huihui Liang, Zheng Li, Xiaomin Si, Jinjin Wang, Qian Liu, Wenyao Xu, Cuicui Yang, Ru-Feng Song, Junrong Li, Tiantian He, Jingyao Li, Xiaoyu Zeng, Jingge Liang, Fang Zhang, Xiaolong Qiu, Yuanyuan Li, Tiantian Bu, Wen-Cheng Liu, Yusheng Zhao, Jinling Huang, Yun Zhou, Chun-Peng Song

## Abstract

Although horizontal gene transfer (HGT) often facilitates environmental adaptation of recipient organisms, whether and how they might affect crop evolution and domestication is unclear. Here we show that three genes encoding cold shock proteins (*CSPs*) were transferred from bacteria to the last common ancestor of Triticeae, a tribe of the grass family that includes several major staple crops such as wheat, barley, and rye. The acquired *CSP* genes in wheat (*TaCSPs*) are functionally conserved with their bacterial homologs by encoding a nucleic acid binding protein. Experimental evidence indicates that *TaCSP* genes positively regulate drought response and improve photosynthetic efficiency under water deficit conditions, by directly targeting a type 1 metallothionein gene to increase ROS scavenging, which in turn contributed to the geographic expansion of wheat. We identified an elite *CSP* haplotype in *Aegilops-tauschii*, introduction of which to wheat significantly increased drought tolerance, photosynthetic efficiency and grain yields. These findings not only provide major insights into the role of HGT in crop adaptation and domestication, but also demonstrate that novel microbial genes introduced through HGT offer a stable and naturally optimized resource for transgenic crop breeding and improvement.

## Introduction

Horizontal gene transfer (HGT) spreads genetic material between distantly related species. Because transferred genes often encode novel functions, HGT may confer adaptive advantages to recipient organisms, allowing them to explore new niches or resources. Although HGT is commonly considered to be a driving force in prokaryotic evolution ^1,2^, recent studies also suggest that HGT has occurred in all major groups of eukaryotes ^3,4^, including complex multicellular eukaryotes such as animals and plants ^5–7^. Particularly, the evolution of plants has been shaped by a considerable number of genes acquired from environmental microbes ^7–9^. These acquired microbial genes not only facilitated plant colonization of land, but also have affected numerous physiological and developmental processes, such as stress response, phytohormone biosynthesis and signaling, vascular development, and secondary metabolism ^7,10–12^. In addition, relatively recent HGT events occurs regularly in nonvascular and seedless vascular plants. Intriguingly, the scale of HGT declines rapidly in seed plants ^7^, and HGT of prokaryotic genes to major crops have not been documented thus far.

Wheat (*Triticum aestivum* L.) is one of the most important staple crops in the world and, along with its close relatives in the tribe Triticeae (the wheat tribe) of the grass family, is mainly distributed in arid and semi-arid areas ^13,14^. Drought stress is a limiting factor for wheat growth and often results in significant grain yield reduction and quality decline ^15,16^. As an allohexaploid, wheat contains three sub-genomes (2*n* = 6*x* =42, AABBDD), showing stronger adaptability and broader geographic distribution than its diploid and tetraploid progenitors ^17–20^. In this work, we report that genes encoding cold shock proteins (CSPs) were horizontally transferred from bacteria to Triticeae and that they are responsive to various abiotic stresses in wheat. Our comprehensive experimental data show that one of the acquired *CSP* genes from the D sub-genome significantly improves drought tolerance and photosynthetic efficiency of wheat under water deficit, through a reactive oxygen species (ROS) scavenging pathway mediated by a type 1 metallothionein gene (*TaMT1*). The expression profile of acquired *CSP* genes in natural populations strongly suggests that they have facilitated the geographic expansion of wheat. Furthermore, we demonstrate that a *CSP* haplotype of enhanced drought tolerance identified in *Aegilops tauschii*, the diploid progenitor of the D sub-genome, could be utilized to significantly increase the survival rate and grain yield of wheat under drought conditions. Our findings provide not only major insights into the role of HGT in crop domestication and adaptation, but also a novel target for transgenic breeding and improvement of wheat and other major crops.

## Results

### Bacterial *CSP* genes were horizontally transferred to Triticeae

Cold shock proteins are characterized by the presence of the cold shock domain (CSD, PF00313), which is about 70 amino acids (aa) in length and widely distributed in both prokaryotes and eukaryotes ^21–24^. In bacteria, these CSPs typically consist of a single CSD domain (CSP-I hereafter) and are believed to be involved in response to multiple stresses ^22,25^. In comparison, eukaryotic CSPs usually include an additional C-terminus (CSP-II hereafter) ^22,26^; in plants, this C-terminus is characterized by the presence of multiple zinc finger motifs (Fig. 1a) ^27^.

**Figure 1.**
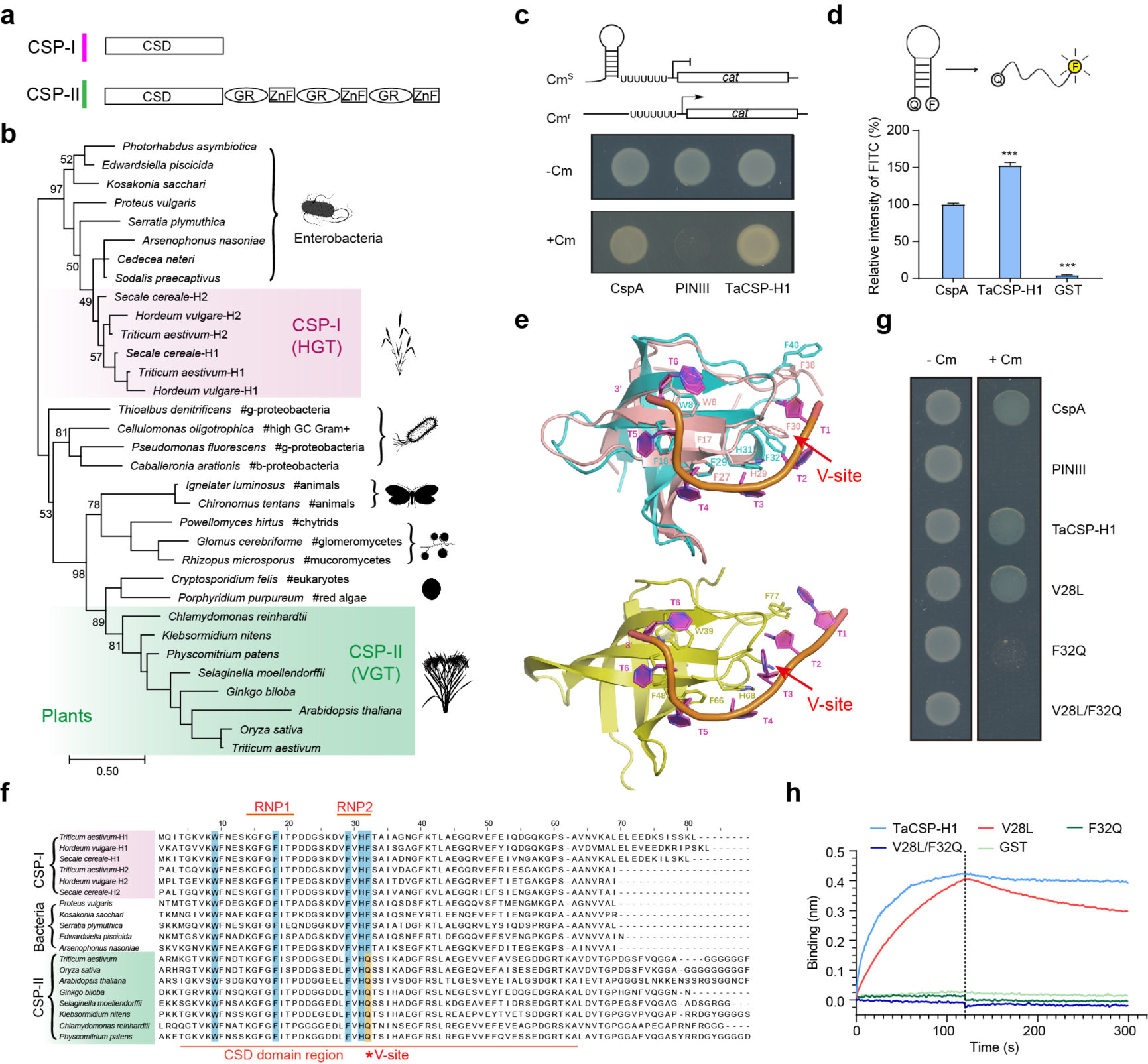
Bacterial *CSP* genes were transferred to Triticeae and encode a nucleic acid binding protein. **a.** Domain and motif organizations in CSP-I and CSP-II proteins. CSD, cold shock domain; GR, glycine-rich region; ZnF, zinc finger. **b.** Phylogenetic analyses of the CSP proteins. Numbers associated with branches show ultrafast bootstrap values from maximum likelihood analyses using IQ-TREE. Bootstrap support values for key branches are noted. **c.** Transcription anti-termination assay of TaCSP-H1 (represented by the homoeolog from D sub-genome unless specified otherwise) in *E. coli* RL211 strain. Overnight liquid cultures of RL211 cells with bacterial CspA, PINIII (empty vector) and TaCSP-H1 were diluted to adjust cell density and spotted onto LB-carbenicillin plates containing 50 μg/ml ampicillin and 1mM IPTG, with or without 15 μg/ml chloramphenicol. **d.** *In vitro* nucleic acid melting activity of TaCSP-H1. Unfolding of the duplex oligodeoxynucleotides relieves quenching and increases the fluorescence intensity. Q, quencher; F, fluorophore. The fluorescence of CspA is set to 100%. Each bar shows mean ± SEM of six replicates (\*\*\**p* < 0.001, Student’s *t*-test). **e.** Structural comparisons between CSPs from different species. Upper: superimposition of Alphafold-predicted TaCSP-H1 (cyan) and (dT)6 bounded BcCspB (*Bacillus caldolyticus* CSP, PDB 2HAX, salmon), both of which are members of CSP-I class. Side chains of highly conserved aromatic residues interacting with the nucleic acid are indicated in stick representation. Lower: cartoon representation of (dT)7 bounded animal CSP LIN28B (*Xenopus tropicalis*, PDB 4A76, yellow), a member of CSP-II class. Only the CSD domain of the protein is used in the analyses. The aromatic side chains that are involved in direct stacking interactions with the (dT)7 bases are shown in stick representation. The V-site of each CSP is indicated by a red arrow. **f.** Sequence alignment of the CSD domain between CSP-I and CSP-II proteins. The positions of RNP1 and RNP2 are indicated by red lines. Conserved aromatic amino acid residues are highlighted, and the red asterisk indicates the V-site of CSPs. **g.** Transcriptional anti-termination assay of the TaCSP-H1 site mutations in *E. coli* RL211. Each of the TaCSP-H1 protein site mutations was transformed into the RL211 strain grown on LB plates with (+Cm) or without (-Cm) chloramphenicol. **h.** Characterization of the nucleic acid binding activities of TaCSP-H1 protein with different site mutations using the BLI assay.

In our genome analyses of wheat, we identified three *CSP-I* genes (eight copies in three sub-genomes), in addition to the eukaryotic *CSP-II* genes (Extended Data Table S1). To assess the distribution of *CSP-I* genes in plants, we performed comprehensive BLAST searches of the NCBI non-redundant (nr) protein sequences, our internally generated genome sequencing data and other related databases, using CSP-I protein sequences of wheat as query. Our results indicated that *CSP-I* genes in land plants are restricted to Triticeae (Extended Data Table S2). Phylogenetic analyses showed that all CSP-I sequences from Triticeae are distantly related to CSP-II homologs but, instead, are closely related to bacterial CSP-I sequences (Fig. 1b, Extended Data Fig. 1). Two of the three CSP-I proteins from Triticeae formed a well-supported monophyletic group within enterobacterial sequences (Fig. 1b), but their relationships with the other CSP-I protein of Triticeae was less certain, largely because of the short sequence length and lack of sufficient resolution. Overall, the above evidence suggests that *CSP-I* genes were transferred from bacteria to the last common ancestor of Triticeae.

We here designate the three horizontally acquired *CSP-I* genes in wheat as *TaCSP-H1*, *TaCSP-H2*, and *TaCSP-H3*, respectively, and the native and vertically inherited *CSP-II* genes as *TaCSP-V*. Our genome analyses indicated that homologs of *TaCSP-H1* and *TaCSP-H2* form a tandem repeat in all published genomes of Triticeae, whereas those of *TaCSP-H3* were only found in species belonging to the B sub-genome (Extended Data Table S2).

### *TaCSP-H1* encodes a nucleic acid-binding protein

Because *TaCSP-H1* and *TaCSP-H2* are tandem duplicates and encode highly similar proteins (82% sequence identity), we selected *TaCSP-H1* for further functional characterization. In bacteria and animals, CSPs typically interact with single-stranded nucleic acids and destabilize their secondary structures ^28,29^. To understand whether TaCSP-H1 possesses similar properties, complementation assays were first performed to assess whether TaCSP-H1 could rescue growth defects of *Escherichia coli* BX04, a mutant strain lacking *CspA/CspB/CspE*/*CspG*, at low temperatures ^30^. As a result, we observed a similar growth phenotype for *E. coli* BX04 expressing *CspA* or *TaCSP-H1* (Extended Data Fig. 2a). Additional transcriptional anti-termination assays in *E. coli* RL211 indicated that both TaCSP-H1 and CspA were able to unfold secondary structures of RNA *in vivo* and, consequently, promote the transcription of chloramphenicol resistance gene *cat* (Fig. 1c). Similarly, both TaCSP-H1 and CspA could directly unfold secondary structures of RNA *in vitro,* under a fluorescein isothiocyanate (FITC) fluorescence molecular beacon system (Fig. 1d). Altogether, these findings suggest that TaCSP-H1 is functionally equivalent to bacterial CspA in nucleic acid binding and unfolding.

### The V-site of TaCSP-H1 is crucial for its nucleic acid melting function

In bacterial CSPs, the cold shock domain consists of two RNA binding motifs (RNP1 and RNP2) ^31^, where sidechains of several aromatic residues are crucial for nucleic acid binding through stacking interactions ^32^. We compared the CSD region between CSP-I and CSP-II proteins and found that, while the first four aromatic residues are highly conserved between the two classes, the fifth (i.e., Phe32 of TaCSP-H1 from the D sub-genome, named V-site hereafter) in CSP-I RNP2 is replaced by Gln (Q) in all sampled CSP-II proteins of plants. In addition, another highly conserved residue in RNP2, Val28 (V28), is also replaced by Leu (L) in CSP-II (Fig. 1f).

We performed transcriptional anti-termination assays to assess whether the F/Q and V/L substitutions affect the nucleic acid binding ability of CSPs. Site mutations at V28 and F32 were generated in *TaCSP-H1*, which in turn was transformed into *E. coli* RL211 on LB plates with (+Cm) or without (-Cm) chloramphenicol. We found that the F32 mutation abolished the nucleic acid unfolding activity of TaCSP-H1, whereas the V28 mutation had no effects (Fig. 1g). In addition, bio-layer interferometry (BLI) assays indicated that the F/Q substitution significantly reduced the substrate binding activity of TaCSP-H1 *in vitro* (Fig. 1h). The above evidence suggests that F32 in RNP2 is essential for TaCSP-H1 in binding nucleic acids and unfolding their secondary structures.

To further understand the effect of V-site, we compared the predicted tertiary structures of TaCSP-H1 and BcCspB (*Bacillus caldolyticus*, PDB 2HAX) ^33^, both of which are members of CSP-I class. In addition, we also simulated the nucleic acid binding using the N-terminal CSD region of LIN28B (*Xenopus tropicalis*, PDB 4A76) ^34^, a CSP-II protein with F/Q substitution at the V-site. As a result, we found the F/Q substitution in LIN28B compromised the aromatic stacking interactions of CSD region to the nucleic acids (Fig. 1e). Because plant CSP-II proteins typically contain several CCHC zinc fingers in the C-terminal, we hypothesized that such compromised nucleic acid affinity might be compensated by the C-terminal zinc fingers. To test this hypothesis, nucleic acid melting activities of the TaCSP-V were assessed, both with and without the zinc fingers. We observed similar nucleic acid folding activities between TaCSP-V and bacterial CspA, but no activity in cells transformed with truncated TaCSP-V that has CSD region alone (Extended Data Fig. 2b-d). This finding is consistent with earlier studies ^35^, suggesting that the C-terminus of TaCSP-V is indispensable for the nucleic acid melting function.

### *TaCSP-H1* positively regulates drought response

To understand how *TaCSP-H* genes affect the physiological activities of wheat, we firstly assessed the expression patterns of *TaCSP-H* homoeologs in different wheat tissues. Analyses of available transcriptomic data showed that *TaCSP-H* homoeologs were mainly expressed in roots, leaves, and spikes (Extended Data Fig. 3a). Because *CSPs* are known to be involved in various stress responses ^36–38^, we again evaluated the expression levels of *TaCSP-H* homoeologs under different stresses (Extended Data Table S3). We found that *TaCSP-H1* and *TaCSP-H2* homoeologs in the D sub-genome responded to various abiotic stresses such as cold, drought, heat, and salt (Fig. 2a, Extended Data Fig. 3b-d), consistent with the diverse abiotic stress responsive *cis*-elements (e.g., AREB, DREB, and STRE) identified in their promoter regions (Extended Data Fig. 4a) ^39,40^. Notably, the expression levels of *TaCSP-H1* (represented hereafter by the copy from D sub-genome unless specified otherwise) homoeologs were not only much higher compared to *TaCSP-H2,* but also significantly induced upon drought treatment (Fig. 2b, Extended Data Fig. 4b-d).

**Figure 2.**
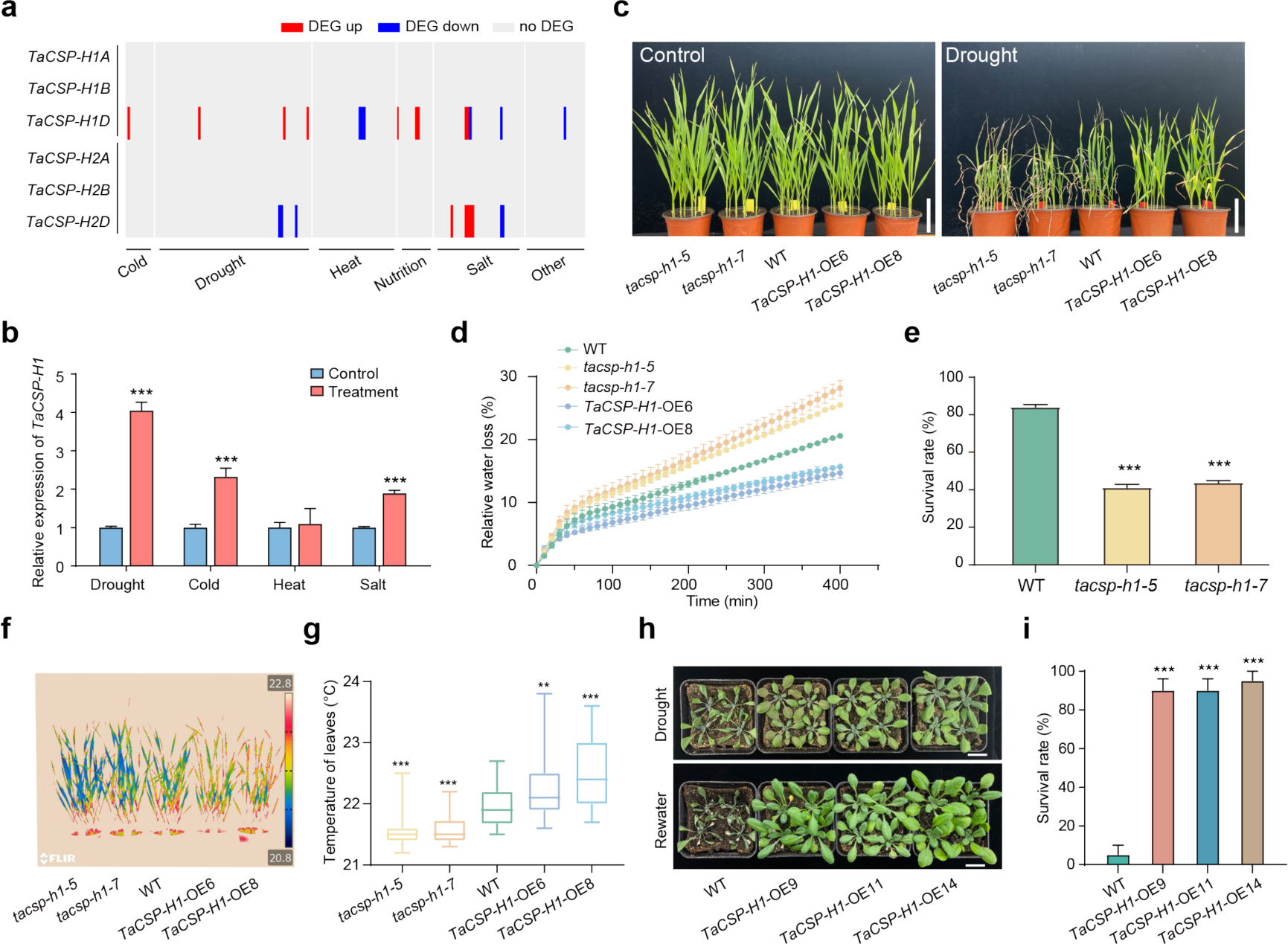
*TaCSP-H1* is involved in abiotic stress responses of wheat. **a.** Differential expression analysis of *TaCSP-H* homoeologs in public transcriptome data. Red and blue rectangles indicate up-regulation and down-regulation, respectively, compared to the control, whereas gray rectangles indicate no significant difference. The differential criteria are set as |Fold Change| > 2 and *q*-value < 0.05. **b.** Expression levels of total *TaCSP-H1* homoeologs in wild-type (WT) seedlings under drought, cold, heat and salt stresses. *TaActin* was used as the internal control. Each bar shows mean ± SEM of three biological replicates (\*\*\**p* < 0.001, Student’s *t*-test). **c.** Assessment of drought tolerance of WT, *tacsp-h1* mutants, and *TaCSP-H1-*OE lines. Wheat seedlings were grown under well-watered conditions for two weeks, and water was then withheld for one week before plants were photographed. Bars = 10 cm. **d.** Relative water loss in WT, *tacsp-h1* mutants, and *TaCSP-H1-*OE lines. Detached leaves of four-week-old plants were used for water loss measurements. Leaves were cut from plants and placed on a piece of weighing paper and weighed every 10 min. Each bar shows mean ± SEM of three replicates, and four leaves from one pot were measured per replicate. **e.** Statistical analysis of survival rates for WT and *tacsp-h1* mutants. Each bar shows mean ± SEM from at least three biological replicates (\*\*\**p* < 0.001, Student’s *t*-test). **f.** Leaf temperatures in WT, *tacsp-h1* mutants, and *TaCSP-H1-*OE lines. Seedlings were grown under normal conditions for two weeks, and water was withheld for two days before measurement. **g.** Statistical analysis of leaf temperatures in (**f**). Ten leaves from one pot were measured per replicate, and three different pots were analyzed (\*\**p* < 0.01, \*\*\**p* < 0.001, Student’s *t*-test). **h.** Soil-grown *TaCSP-H1*-OE lines of *Arabidopsis* seedlings are more drought tolerant than the WT. Twenty-one-day-old seedlings were subjected to drought stress by withholding water for 14 days. The plants were photographed before and after rewatering. Bars = 2 cm. **i.** Statistical analysis of survival rates for WT and *TaCSP-H1*-OE lines in (**h**). Each bar represents mean ± SEM from at least 20 seedlings.

We knocked out the *TaCSP-H1* homoeologs from all three sub-genomes simultaneously in the wheat *cultivar* Fielder, using CRISPR/Cas9 technology, and generated two homozygous mutants, designated here as *tacsp-h1-5* and *tacsp-h1-7* (Extended Data Fig. 5). Under normal conditions, the mutants showed no visible difference compared with the wild-type (WT) plants during their entire life cycle (Extended Data Fig. 6). However, the *tacsp-h1* mutants exhibited defects to varying degrees under various stress treatments (Extended Data Fig. 7a-c). Particularly, the *tacsp-h1* mutants became visibly wilted two days after drought stress treatment and had an about 50% lower survival rate than WT (Fig. 2e, Extended Data Fig. 7d). We further assessed other drought-related phenotypes in the *tacsp-h1* mutants and *TaCSP-H1* overexpression (*TaCSP-H1*-OE) lines. Compared with WT, the *tacsp-h1* mutants had an ∼30% increase in water loss rate (Fig. 2d) and ∼0.5℃ lower seedling leaf temperature (Fig. 2f-g), indicating much higher transpiration strength of the *tacsp-h1* mutants. In comparison, the *TaCSP-H1*-OE lines exhibited drought resistant phenotypes compared with WT (Fig. 2c-d, f-g), and ectopic expression of *TaCSP-H1* in *Arabidopsis* improved plant drought tolerance (Fig. 2h-i, Extended Data Fig. 7e). Collectively, these results indicate that *TaCSP-H1* is a positive regulator of drought response in wheat.

### *TaCSP-H1* contributes to the spread of wheat by improving photosynthetic efficiency

To investigate how *TaCSP-H1* regulates drought response in wheat, we generated RNA-seq data using WT, *tacsp-h1*, and *TaCSP-H1-*OE. As a result, numerous differentially expressed genes (DEGs) were identified before and after drought treatments (Fig. 3a, Extended Data Fig. 8a-b, Extended Data Table S5). Notably, most of the identified DEGs are enriched in “photosynthesis”, “tetrapyrrole metabolism”, “chlorophyll metabolic processes”, and “response to oxygen radicals” (Extended Data Fig. 8c, Extended Data Table S6), many of which showed opposite expression patterns between *tacsp-h1* and *TaCSP-H1-*OE (Extended Data Fig. 8d). As such, we assessed the photosynthesis-related characters in WT, *tacsp-h1*, and *TaCSP-H1-*OE. Compared with WT, both the photosynthetic rate and the photosystem II efficiency increased significantly in *TaCSP-H1*-OE but decreased in *tacsp-h1* mutants under drought conditions (Fig. 3b, Extended Data Fig. 9a), indicating that the expression level of *TaCSP-H1* is positively correlated with photosynthetic efficiency. Because photosynthetic efficiency is a key trait for crop adaptation and distribution ^41^, we further analyzed the expression patterns of *TaCSP-H1* in natural populations of wheat. Overall, the expression levels of *TaCSP-H1* were lower in populations among the western coast of the Caspian Sea and in the Fertile Crescent, where wheat originated ^17,42–44^, and became increasingly higher in populations of other expanded geographic areas (Extended Data Fig. 9b-c) ^45^. The above evidence suggests that *TaCSP-H1* has facilitated the adaptation and geographic expansion of wheat across the world.

**Figure 3.**
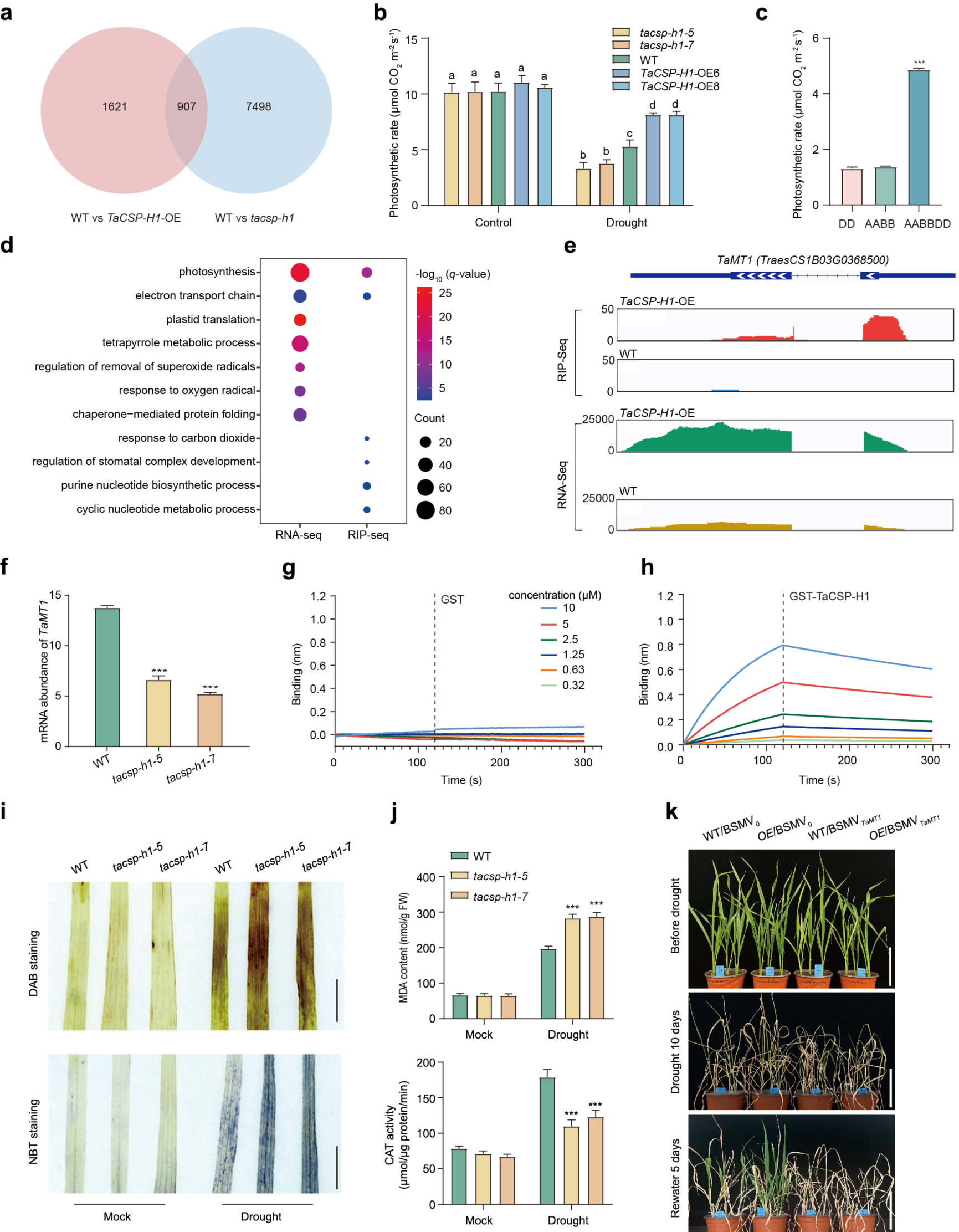
*TaCSP-H1* improves photosynthetic efficiency of wheat and promotes ROS scavenging via *TaMT1* under drought stress. **a.** Venn diagram of total numbers of DEGs in WT vs *TaCSP-H1*-OE and WT vs *tacsp-h1* comparisons under drought conditions. **b.** Photosynthetic rates of flag leaves in WT, *tacsp-h1* mutants, and *TaCSP-H1-*OE lines. Each bar shows mean ± SEM of six biological replicates. Different letters represent significant differences (*p* < 0.05, one-way ANOVA, Tukey’s HSD test). **c.** Photosynthetic rates of flag leaves in diploid wheat (*Ae. tauschii*, 2*n* = 14, DD), tetraploid wheat (*T. durum*, 2*n* = 4*x* = 28, AABB) and synthetic hexaploid wheat (2*n* = 6*x* = 42, AABBDD). The synthetic hexaploid wheat was generated by hybridization of diploid and tetraploid wheat, followed by artificial doubling at F1 generation. Each bar shows mean ± SEM of six biological replicates (\*\*\**p* < 0.001, Student’s *t*-test). **d.** GO enrichment analysis of RNA-seq and RIP-seq data. DEGs between WT and *TaCSP-H1-* OE under normal conditions were used as RNA-seq data set. The differential genes between WT-IP and *TaCSP-H1*-OE-IP were used as RIP-seq data set. Each circle represents a GO term, and the number of genes enriched in a pathway corresponds to the size of the circle. The degree of significance for the enrichment of DEGs in a pathway is represented by -log_10_ (*q*-value). **e.** Profile of TaCSP-H1 binding to the transcripts of *TaMT1* and the abundance of *TaMT1* transcript in WT and *TaCSP-H1-*OE. The genomic structure of *TaMT1* is shown at the top. The dotted line indicates the intron of *TaMT1*. Wiggle plots of RIP-seq and RNA-seq are shown below the gene structure. The y-axis indicates sequencing read abundance of RIP-seq or RNA-seq data. The x-axis indicates the chromosomal position of *TaMT1* gene. **f.** The mRNA abundance of *TaMT1* decreases in *tacsp-h1* mutants. The primers were designed at the second exon and the 3’-UTR of *TaMT1. TaActin* was used as the internal control. Each bar shows mean ± SEM of three biological replicates (\*\*\**p* < 0.001, Student’s *t*-test). **g-h**. Characterization of the binding of TaCSP-H1 protein at increasing concentrations to *TaMT1* mRNA using the BLI assay. GST (**g**) or GST-TaCSP-H1 (**h**) proteins were incubated with the synthesized biotin-labeled RNA probes, and the probes were designed at the first exon of *TaMT1*. **i.** ROS levels revealed by DAB and NBT staining in leaves of wheat seedlings under well-watered and water-withheld conditions. Bars = 1 cm. **j.** Malonaldehyde (MDA) contents and catalase (CAT) activities in leaves of WT and *tacsp-h1* mutants under normal and drought conditions. Each bar shows mean ± SEM of six biological replicates (\*\*\**p* < 0.001, Student’s *t*-test). **k.** Silence of *TaMT1* abolishes the role of *TaCSP-H1* in drought tolerance. WT/BSMV_0_: WT transformed with empty VIGS vector; OE/BSMV_0_: *TaCSP-H1-*OE line transformed with empty VIGS vector; WT/BSMV*_MT1_*: WT transformed with *TaMT1* VIGS vector; OE/BSMV*_MT1_*: *TaCSP-H1-*OE line transformed with *TaMT1* VIGS vector. Bars = 10 cm.

Compared with its diploid and tetraploid ancestors *Ae. tauschii* (2*n* = 14, DD) and *T. durum* (2*n* = 4*x* = 28, AABB), wheat exhibits stronger environmental adaptation and wider distribution ^17,20^. We therefore speculated that this increased adaptability of wheat might be associated with elevated photosynthetic efficiency and *CSP-H1* expression. We tested this hypothesis using a synthetic hexaploid wheat line so that its direct diploid and tetraploid parents could be unambiguously determined. We found that the photosynthetic rate of synthetic hexaploid wheat was about 3-fold higher compared with its diploid and tetraploid parents (Fig. 3c). As expected, comparative transcriptomic analyses between the synthetic wheat and its diploid and tetraploid parents revealed that many photosynthesis-related genes were differentially expressed among them (Extended Data Fig. 10a, Extended Data Table S7-S8), some of which were also observed in the comparison of *TaCSP-H1*-OE and WT (Extended Data Fig. 10b). Importantly, the expression level of *CSP-H1* homoeologs increased from diploid to hexaploid wheat, especially under drought stress (Extended Data Fig. 10c). The above data suggest that the increasing photosynthetic efficiency during the origin of hexaploid wheat was likely associated with a higher level of *TaCSP-H1* expression.

### TaCSP-H1 directly binds to the *TaMT1* transcript and promotes ROS scavenging

Given that TaCSP-H1 can directly bind to RNAs and unfold their secondary structures (Fig. 1c-d), we also performed RNA immunoprecipitation sequencing (RIP-seq) to search for its possible direct targets (Extended Data Table S9). Most genes identified in RIP-seq data are related to “photosynthesis”, “response to carbon dioxide”, or “regulation of stomatal complex development” (Fig. 3d, Extended Data Table S10), again supporting the role of *TaCSP-H1* in drought responses and photosynthesis. We identified 35 overlapped genes between the RNA-seq and RIP-seq datasets (Extended Data Table S11) and validated a subset of them by quantitative real-time PCR (Extended Data Fig. 11). Notably among the overlapped genes, the expression level of *TaMT1* (*TrasesCS1B03G0368500*), a type 1 metallothionein reportedly involved in ROS scavenging in rice ^46^, was significantly increased in *TaCSP-H1-*OE (Fig. 3e). In comparison, levels of both *TaMT1* transcript and its encoded protein decreased significantly in *tacsp-h1* mutants (Fig. 3f, Extended Data Fig. 12). This evidence suggests that *TaMT1* is likely a downstream target of *TaCSP-H1* in regulating drought tolerance. Indeed, our BLI assays using GST-TaCSP-H1 fused protein found that TaCSP-H1 bound strongly to the *TaMT1* transcript in a concentration-dependent manner (Fig. 3g-h). Altogether, the above results demonstrate that TaCSP-H1 promotes *TaMT1* expression by directly binding to its transcript.

Because the ortholog of *TaMT1* regulates drought response in rice by reducing ROS accumulation ^46^, we decided to analyze the ROS level in *tacsp-h1* mutants using diaminobenzidine (DAB) and nitroblue tetrazolium (NBT) staining. The ROS levels were comparable between WT and *tacsp-h1* mutants under normal growth conditions. However, ROS level in the *tacsp-h1* mutants was much higher compared to WT under drought treatment, (Fig. 3i). Meanwhile, the malonaldehyde (MDA) content increased ∼40 % in *tacsp-h1* mutants whereas catalase activities decreased ∼35 % (Fig. 3j), suggesting that TaCSP-H1 reduces the ROS level under water deficit conditions. We further tested whether *TaMT1* is required for *TaCSP-H1*-mediated drought tolerance using the virus induced transient gene silencing (VIGS) in WT and *TaCSP-H1*-OE plants. As a result, *TaMT1* knock-down not only caused a drought sensitive phenotype in WT, but also rendered the *TaCSP-H1*-OE plants sensitive to drought stress (Fig. 3k), indicating that *TaMT1* is indeed required for *TaCSP-H1* to regulate drought response. Collectively, these findings show that *TaCSP-H1* positively regulates drought tolerance through a *TaMT1*-mediated ROS scavenging mechanism under drought stress.

### Natural variation of *CSP-H1* in *Ae. tauschii* provides an elite allele for wheat breeding

Because *TaCSP-H1* in the D sub-genome is the most expressed gene and mightily induced by drought stress (Fig. 2a, Extended Data Fig. 4), we searched the available sequencing data of the D sub-genome and its progenitor *Ae. tauschii* to identify beneficial *CSP-H1* variations for wheat improvement. Overall, we found very limited allelic variation in the D sub-genome but, instead, tremendous variation in *Ae. tauschii* (Extended Data Fig. 13). Specifically, an indel in the coding sequence of *CSP-H1* was identified in *Ae. tauschii*, which divided *CSP-H1* into two haplotypes, named here *AetHap1* and *AetHap2*, respectively (Fig. 4a). Notably, *AetHap1* encodes a protein identical to TaCSP-H1 from the D sub-genome, consistent with the earlier findings that the D sub-genome of wheat evolved only 8,000-9,000 years ago from *Ae. tauschii* ^20,44,47^. The two haplotypes of *Ae. tauschii* have different distribution patterns – while *AetHap1* is largely restricted to the northern coast of the Caspian Sea and the Caucasus, *AetHap2* is widely distributed from the Caucasus through Central Asia to Central China (Fig. 4b, Extended Data Table S12). Because water deficit often limits plant distribution and *TaCSP-H1* positively regulates drought response (Fig 2d-i), we speculated that *AetHap1* and *AetHap2* may be associated with different levels of drought tolerance. We tested this hypothesis using drought tolerance index (DTI) and, indeed, found that *Aegilops* plants carrying *AeHap2* had stronger drought tolerance than those carrying *AetHap1* (Fig. 4c, Extended Data Fig. 14).

**Figure 4.**
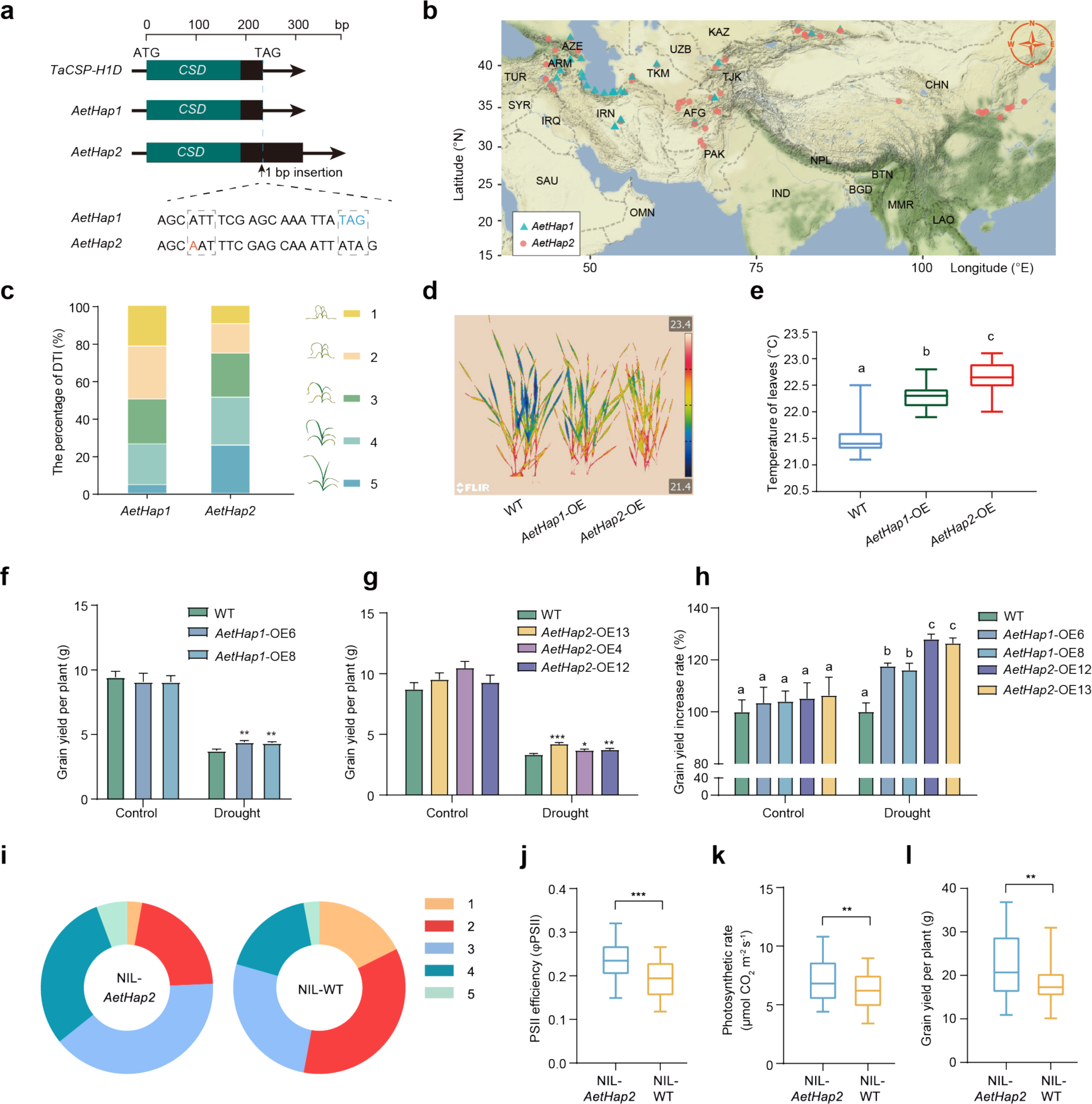
*AetCSP-H1* natural variations improve wheat drought adaptation and grain yield. **a.** Genomic sequences of *TaCSP-H1D* and two *AetCSP-H1* haplotypes (*AetHap1* and *AetHap2*) in *Ae. tauschii*. *AetHap2* contains a base pair insertion in the coding sequence at 226 bp from the initiation codon and generates a frameshift at stop codon. 52 *AetHap1* accessions and 133 *AetHap2* accessions were used for statistical analyses. **b.** Geographic distribution of the two *AetCSP-H1* haplotypes. Blue triangles and red circles indicate *AetHap1* and *AetHap2*, respectively. **c.** Percentage distribution of drought tolerance index (DTI) in the two *AetCSP-H1* haplotypes from *Ae. tauschii*. 46 *AetHap1* accessions and 51 *AetHap2* accessions were used for statistical analyses. **d.** Leaf temperatures in WT and *AetHap1-*OE or *AetHap2-*OE lines. Seedlings were grown under normal conditions for two weeks, and water was withheld for two days before measurement. **e.** Statistical analysis of leaf temperatures in (**d**). Ten leaves from one pot were measured per replicate, and three different pots were analyzed. Different letters represent significant differences (*p* < 0.05, one-way ANOVA, Tukey’s HSD test). **f.** Grain yields of WT and *AetHap1-*OE lines under well-watered and water-deficit conditions (n = 20 biologically independent plants; \*\**p* < 0.01, Student’s *t*-test). **g.** Grain yields of WT and *AetHap2-*OE lines under well-watered and water-deficit conditions (n = 15 biologically independent plants; \**p* < 0.05, \*\**p* < 0.01, \*\*\**p* < 0.001, Student’s *t*-test). **h.** Grain yield increase rates of *AetHap1-*OE and *AetHap2-*OE lines under well-watered and water-deficit conditions. The grain yields of WT under normal and drought conditions were set as 100%, respectively. Different letters represent significant differences (*p* < 0.05, one-way ANOVA, Tukey’s HSD test). **i.** Percentage distribution of drought tolerance index (DTI) of wheat near isogenic lines (NILs) harboring *AetHap2* and *TaCSP-H1D* (WT). The introgression lines used for statistics were 70 (NIL-*AetHap2*) and 68 (NIL-WT), respectively. **j.** Photosystem II (PSII) efficiency (φPSII) of flag leaves in NILs harboring *AetHap2* or *TaCSP-H1D*. The introgression lines used for statistics were 60 (NIL-*AetHap2*) and 60 (NIL-WT), respectively (\*\*\**p* < 0.001, Student’s *t*-test). **k.** Photosynthetic rate of flag leaves in NILs harboring *AetHap2* or *TaCSP-H1D*. The introgression lines used for statistics were 60 (NIL-*AetHap2*) and 60 (NIL-WT), respectively (\*\**p* < 0.01, Student’s *t*-test). **l.** Grain yield per plant of NILs harboring *AetHap2* or *TaCSP-H1D*. The introgression lines used for statistics were 60 (NIL-*AetHap2*) and 60 (NIL-WT), respectively (\*\**p* < 0.01, Student’s *t*-test).

To assess the potential of *AetHap2* in wheat improvement, we produced overexpression lines of *AetHap1* (i.e., *TaCSP-H1* from the D sub-genome of wheat) and *AetHap2* in wheat. Water loss assays using detached leaves showed that wheat overexpressing *AetHap2* (*AetHap2*-OE) had lower water loss than both WT and that overexpressing *AetHap1* (*AetHap1*-OE) (Extended Data Fig. 15a). Meanwhile, leaf temperatures of *AetHap2*-OE were higher than WT and *AetHap1-*OE (Fig. 4d-e), indicating that *AetHap2* could improve wheat drought tolerance. Additional BLI and complementation assays indicated that AetHap2 had ∼13.8-fold stronger nucleic acid binding and higher melting activities than AetHap1 (Extended Data Fig. 15c-d), suggesting that *AetHap2* improves drought tolerance through an enhanced nucleic acid binding capability, consistent with the significant up-regulation of *TaMT1* under drought conditions (Extended Data Fig. 15b). We further investigated the effects of *AetHap2* on agronomic traits. Overexpressing of *AetHap1* and *AetHap2* both could improve the grain yield compared to WT under water-deficit conditions (Fig. 4f-g), and the grain yield of *AetHap2*-OE was ∼10% higher than that of *AetHap1*-OE (Fig. 4h). Taken together, these data demonstrate that *AetHap2* has a greater potential than *AetHap1* in improving drought tolerance and grain yield of wheat.

Because *AetHap2* is absent in the wheat genome, we also introduced it into a modern wheat cultivar through introgression. An accession KS262 harboring *AetHap2* was identified from introgression lines of T093 (an accession of *Ae. tauschii* carrying *AetHap2*) and an elite wheat cultivar Zhoumai18 ^47^ (Extended Data Fig. 16), which in turn was backcrossed with Zhoumai18 to create *TaCSP-H1* near isogenic lines (NILs) (Extended Data Fig. 17a). We then randomly selected 70 individuals carrying *AetHap2* (NIL-*AetHap2*) and 68 individuals carrying *TaCSP-H1* (NIL-WT) using the Kompetitive Allele-Specific PCR (KASP) markers for comparative drought tolerance assays (Extended Data Fig. 17b). Overall, we found that introgression individuals carrying *AetHap2* were significantly more drought tolerant (Fig. 4i), and their photosystem II efficiency and photosynthetic rate were much higher (Fig. 4j-k). Meanwhile, the grain yield per plant had an ∼20% increase with the introgression of *AetHap2* segments (Fig. 4l). These data again suggest that *AetHap2* is an elite allelic variation for modern wheat improvement.

### *TaCSP-H1* offers a novel resource for major crop improvement

Given the absence of *TaCSP-H1* from many other crops, we decided to test their applications in transgenic improvement of other crops. *TaCSP-H1* was transferred into rice to assess its effects on drought tolerance and grain yield. As a result, we found that overexpressing *TaCSP-H1* could improve drought tolerance and seedling survival rate in rice (Extended Data Fig. 18). Compared with WT, the *TaCSP-H1*-OE lines under water deficit conditions exhibited significantly greater drought tolerance at heading stage (Fig. 5a) and higher grain yields (Fig. 5b-c). In addition, transcriptomic data revealed that *TaCSP-H1* overexpression altered photosynthesis and diverse abiotic stress signaling pathways in rice (Extended Data Table S13-S14), especially the response to water deprivation (Fig. 5d). These data demonstrate that *TaCSP-H1* is functionally conserved in drought tolerance in rice and provides a novel candidate for the breeding and improvement of other crops.

**Figure 5.**
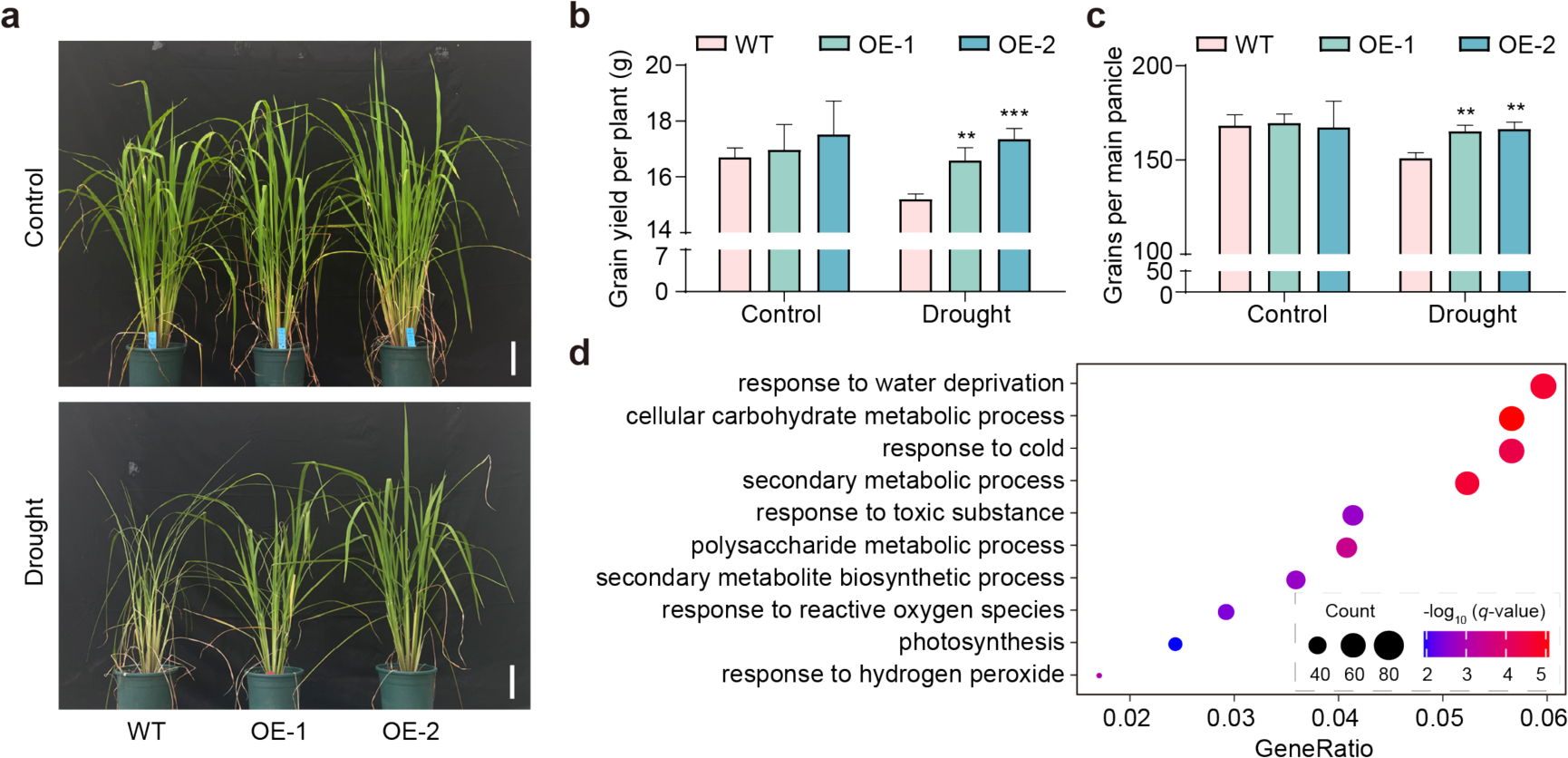
*TaCSP-H1* increases abiotic stress response and grain yield in rice. **a.** Phenotypes of WT and *TaCSP-H1-*OE lines (OE-1 and OE-2) in rice under well-watered and water-deficit conditions. Rice seedlings were grown under well-watered conditions for two months, and water was withheld for two weeks before plants were photographed. Bars = 15 cm. **b-c**. Agronomic traits of WT and *TaCSP-H1*-OE lines (OE-1 and OE-2) under well-watered and water-deficit conditions (n = six biologically independent plants; \*\**p* < 0.01, \*\*\**p* < 0.001, Student’s *t*-test). **d**. GO enrichment analysis of DEGs in WT and *TaCSP-H1*-OE lines under drought conditions. Each circle represents a GO term, and the number of genes enriched in a pathway corresponds to the size of the circle. The abscissa indicates the ratio of the number of DEGs annotated to a particular pathway to the number of the DEGs annotated to all pathways. The degree of significance for the enrichment of DEGs in a pathway is represented by -log_10_ (*q*-value).

## Discussion

Despite the historical controversies around HGT in complex multicellular eukaryotes ^48–50^, an increasing amount of genome data indicate that genes acquired from environmental microbes have played a key role in plant evolution ^5,7–9^. Nonetheless, the distribution of HGT in plants is extremely uneven ^7^. While a considerable number of genes of microbial origin are reported in non-seed plants, very few are known in seed plants, with the notable exception of the grass family. Thus far, several genes of fungal origin were reportedly transferred to different groups of the grass family, all of which presumably involved *Epichloe* endophytes as the donor ^11,51,52^. Unlike these previous reports, *CSP-H* genes reported here were acquired from free-living bacteria by the last common ancestor of Triticeae. Together with the previous reports ^11,51,52^, our findings suggest that HGTs in the grass family are active and dynamic, and they can be derived from various other microbial sources (Fig.1b, Extended Data Fig. 1a). Given the public concerns of biosafety related to transgenic crops ^53–55^, why and how the grass family is receptive to foreign genes merit further investigations.

The finding that *TaCSP-H1* promotes both drought tolerance and photosynthesis has major implications in understanding the evolution and adaptation of wheat and related crops. Drought induces stomatal closure and reduces CO_2_ uptake, ultimately leading to lower photosynthetic efficiency and accumulation of ROS ^56–58^. By directly targeting *TaMT1*, *TaCSP-H1* mediates ROS scavenging (Fig. 3g-k), which in turn could alleviate drought stress and improve photosynthesis. The increasing level of *TaCSP-H1* expression from diploid *Ae. tauschii* to hexaploid wheat (Extended Data Fig. 10c), as well as from the coast of the Caspian Sea to other geographic areas (Extended Data Fig. 9c), strongly suggests that *TaCSP-H1* might have not only contributed to the formation of hexaploid wheat by scavenging excessive ROS produced as a by-product of elevated photosynthetic activities, but also facilitated its subsequent geographic expansion. Importantly, the tribe Triticeae includes several other major staple crops such as rye (*Secale cereale*), barley (*Hordeum vulgare*), and their wild relatives, many of which (e.g., *Elymus* spp.) are also of economic importance. Particularly, wheat, barley and rye are among the most widely distributed and cultivated crops in the word ^59^. Although our current study is focused on the role of *TaSCP-H1* from the D sub-genome in drought tolerance, it should be emphasized that wheat contains eight *TaCSP-H* copies, including three *TaCSP-H1* and three *TaCSP-H2* (one from each of the three sub-genomes), plus two *TaCSP-H3* from the B sub-genome. Because the CSD domain is known to be involved in growth and developmental processes, as well as responses to other abiotic stresses ^22,23,29,60^, it is most likely that these *CSP-H* genes participate in various other processes. The likely involvement of *CSP-H* genes in response to other stresses is supported by the transcriptomic data (Fig. 2a) and consistent with the fact that wheat, barley, rye, and their wild relatives from Triticeae are mostly found in arid and harsh environments of temperate zones. Detailed investigations on other *CSP-H* genes in wheat and related species from Triticeae should provide a clearer picture to understand their role in crop adaptation and evolution.

HGT represents a process of natural transgenic engineering, of which the effectiveness was tested and optimized by natural selection over time. Bacterial *CSP* genes were previously overexpressed in several crops (e.g., maize, rice, and wheat) to increase their tolerance to stresses, including drought, and grain yields ^36,61^. Our data show that naturally transferred bacterial *CSP* genes are effective in the enhancement of drought tolerance and photosynthetic efficiency of wheat (Fig. 2c-g, Extended Data Fig. 9). In addition, variation of these naturally transferred genes are further selected under various environmental conditions, thus creating potentials for greater improvement of crops. Our study indicates that *AetHap2* confers significantly stronger drought tolerance and greater grain yields (Fig. 4d-h), suggesting that this is an elite haplotype for breeding and improvement of wheat and other crops. Importantly, microbial genes introduced through HGT and subsequent variations, such as *AetHap2*, are both stable and naturally optimized, providing a theoretical basis for their utilization in modern transgenic engineering and synthetic biology.

## Materials and Methods

### Plant materials and growth conditions

Wheat cv. Fielder was used as the wild type unless specified otherwise. Wheat seeds were germinated on petri dishes for five days and then moved to soil under a 16-h-light/8-h-dark photoperiod, with 60% relative humidity at 22℃. For drought treatments, water was withheld at the five-leaf stage of plant, and soil water content was maintained at 8% until rewatering.

### Vector construction and wheat genetic transformation

To construct *TaCSP-H1* overexpressing vector, the full-length coding sequence (CDS) of *TaCSP-H1D* fused with the 3×Flag tag sequence was cloned into pMWB110 to achieve the *Ubi:TaCSP-H1-Flag* construction. To create gene knockout constructs, target sites were designed to recognize conserved regions of homologous genes using the website E-CRISP Design (http://www.e-crisp.org/E-CRISP/) and then cloned into the CRISPR/Cas9 vector pBUE411 ^62^. All binary vectors harboring the desired constructs were transferred into *Agrobacterium tumefaciens* strain EHA105 and then transformed into wheat cultivar Fielder by *Agrobacterium*-mediated wheat immature embryo transformation method ^63^.

### HGT identification and phylogenetic analysis

HGT candidates in *T. aestivum* were identified by searching the latest NCBI nonredundant (nr) protein sequence database (downloaded on 2023.12.25) using the method described in Ma et al. ^7^. To construct the phylogenetic relationship of *CSP* genes, protein sequences of *TaCSP-H1D* (*TraesCS2D03G0314000*), *TaCSP-H2D* (*TraesCS2D03G0314100*), *TaCSP-H3* (*TraesCS2B02G511500LC*) and *TaCSP-V* (*TraesCS6A02G069500*) of *T. aestivum* were used as query to search against nr database using BLASTP (E-value cutoff = 1e-10). Homologous protein sequences from representative lineages of three domains of life (bacteria, archaea, and eukaryotes) were sampled for phylogenetic analyses. Multiple sequence alignments were performed using Mafft ^64^, with default settings. Gaps and ambiguously aligned sites were removed or trimmed manually. Phylogenetic analyses were performed with a maximum likelihood method using IQ-Tree ^65^. Best-fitting amino acid substitution model was selected based on BIC. Branch support values were estimated with 1,000 replicates using ultrafast bootstrapping in IQ-Tree.

### Thermal imaging

Thermal imaging of wheat seedlings was conducted as described previously with modifications ^66^. Briefly, the seedlings were grown at 22°C under a 16-h-light/8-h-dark cycle in a light chamber for three weeks. Thermal images were obtained using a ThermaCAMSC1000 infrared camera (FLIR). Leaf temperatures were measured using IRBIS 3 professional software. For each genotype, plants in three different pots were analyzed with similar results.

### Protein expression and purification

Protein expression and purification were performed as described previously with modifications ^67^. GST-TaCSP-H1 proteins were expressed in *E. coli* strain BL21 with 0.5 mM isopropyl β-D-1-thiogalactopyranoside (IPTG) for 18 h at 25°C. The culture was then collected by centrifugation at 5000 *g* for 10 min at 4°C. The pellet was resuspended with 1× PBS (150 mM NaCl, 10 mM Na_2_HPO_4_, 2 mM KH_2_PO_4_, 2.7 mM KCl, pH 7.4, 1 mM PMSF, and 1× cocktail) and sonicated for 10 min. After centrifugation for 30 min at 10000 *g*, the supernatant was incubated with glutathione Sepharose 4B (GE) resin for 3 h at 4°C. The resin was then washed three times with 1× PBS to remove nonspecific binding proteins. Finally, the GST-TaCSP-H1 proteins were eluted from the resin with GSH buffer (50 mM Tris-HCl, pH 8.0, and 10 mM GSH).

### RNA extraction, library construction and sequencing

Total RNAs were extracted from 3-week-old seedlings of WT, *tacsp-h1* and *TaCSP-H1*-OE lines for each of three biological replicates using TRIzol (Invitrogen) and treated with DNase I to remove the genomic DNA according to the user manual. The mRNA was isolated and fragmented using mRNA capture beads (VAHTS mRNA Capture Beads, N401). The resultant RNA samples were reverse-transcribed for library construction, using the NEB Next® VAHTS Universal V6 RNA-seq Library Prep Kit for Illumina, NR604. The libraries were purified by DNA clean beads (VAHTS DNA Clean Beads, N411) to select for fragments of 200-300 bp, and analyzed on Agilent 2100 Bioanalyzer (Agilent) to measure fragment size distribution. The qualified libraries were sequenced on Illumina Nova Seq 6000 (Extended Data Table S4).

RNA-seq data were processed with the nf-core RNA-seq pipeline v3.12.0 ^68^. Briefly, the pipeline was based on Nextflow v23.04.1 and processed data using the parameters ‘--extra_star_align_args "--seedPerWindowNmax 20" --trimmer fastp --aligner star_salmon -- bam_csi_index’. Low-quality reads were removed from the raw reads with Fastp ^69^, whereas remaining reads were mapped to the reference genome wheat CSv2.1 ^70^ with STAR v2.7.9a ^71^. Transcript quantification was done with salmon v1.10.1 ^72^. DEGs between control and stress samples were detected using the R procedure in Bioconductor package DESeq2 ^73^. Eggnog-mapper ^74^ was applied to annotate GO, KEGG and gene functions. Enrichment analyses of DEGs in GO terms and KEGG pathways were performed using ClusterProfiler ^75^.

### RNA immunoprecipitation sequencing and analysis

Three-week-old WT and *TaCSP-H1*-OE line seedlings were harvested, ground into powder in liquid nitrogen, and then lysed. Cells were crosslinked in 0.3% formaldehyde solution for 10 min, followed by reaction termination in 125 mM glycine. The cell pellet was washed twice with 1×PBS and then lysed in RIP buffer (25 mM Tris, pH 7.4, 150 mM KCl, 5 mM EDTA, 0.2% CA-630, 0.05% SDS) at 4℃ for one hour. The cell lysate containing nucleic acid-protein complexes was sheared by ultrasonication and centrifuged to obtain the supernatant, one-tenth of which was aliquoted as input. The remaining supernatant was subjected to immunoprecipitation by adding 5 μg anti-Flag antibody conjugated with Dynabeads™ Protein A (Thermo Fisher) and incubated at 4℃ for 16 h. Antibody-Flag-nucleic acid complexes were collected by magnetic beads. Both immunoprecipitated and input samples were digested with DNase I at 37℃ for 15 min to remove DNA contamination, followed by protein removal by digestion with proteinase K at 55℃ for 30 min. The resulting RNA fragments were purified with phenol: chloroform: isoamyl alcohol (25:24:1) and subjected to library construction with the NEB Next® Ultra TM RNA Library Prep Kit for Illumina®. Constructed libraries were purified by purification beads (Vazyme, N411) and analyzed on Agilent 2100 Bioanalyzer (Agilent) to measure fragment size distribution (around 150 bp). Qualified libraries were sequenced on Illumina Nova Seq 6000 (Extended Data Table S4).

RIP-seq data analysis followed previous studies with slight modifications ^76^. Briefly, the pipeline was based on Nextflow v23.04.1 and processed data using the parameters ‘--extra_star_align_args "--seedPerWindowNmax 20" --remove_ribo_rna true --trimmer fastp --aligner star_salmon --bam_csi_index’. Low-quality reads were removed from the raw reads with Fastp ^69^, whereas remaining reads were mapped to the reference genome wheat CSv2.1 ^70^ with STAR v2.7.9a ^71^. Transcript quantification was done with salmon v1.10.1 ^72^. Putative RIP target genes were determined based on common targets identified by two methods. a. Comparison of IP and input read counts using DESeq2 ^73^ (FC > 1.5 and p-value < 0.05). b. Comparison of OE_IP and WT_IP using default parameters in MACS3 ^77^ (p-value < 0.05). Sequence profiles showing the binding sites were visualized using IGV.

### Biolayer Interferometry (BLI) assay

BLI was carried out on a Sartorius Octet system according to the user guide. 5’-biotin-labeled-RNA probes were synthesized from Sangon Biotech. Streptavidin biosensors were hydrated for 10 min prior to the experiment in a buffer solution containing 15 mM HEPES (pH 7.0), 100 mM KCl, and 5 mM MgCl_2_. Biotin-labeled RNA probes were diluted in the same buffer to 1 μM. Protein samples were diluted to different concentrations in the PBS buffer. After loading and quenching, the sensors fixed by labeled RNA probes were incubated with the protein samples to measure the binding affinity. Dissociation constant (Kd) values and standard deviations were generated by BLItz Pro software analysis.

### Quantitative real-time PCR (qRT-PCR) analysis

Total RNA was extracted from wheat plants using TRIzol reagent (Invitrogen). Extracted RNA was reverse transcribed to cDNA using a PrimeScript^TM^ RT reagent Kit (TaKaRa). The qRT-PCR was performed with SYBR Premix Ex Taq^TM^ (TaKaRa) on a LightCycler 480 II (Roche). The wheat gene *TaActin* (*TraesCS5D03G0333200*) was used as an internal reference for normalization of qRT-PCR threshold cycle values. For each sample, at least three biological replicates were performed with similar results. Primer sequences used in the analyses are listed in Extended Data Table S15.

### Virus-induced gene silencing (VIGS)

Barley streak mosaic virus (BSMV) vectors were used to conduct VIGS constructs. The targeted region of *TaMT1* (*TrasesCS1B03G0368500*) was amplified from cDNA of the Fielder wheat cultivar and cloned into the BSMV-γ vector. BSMV viral particles containing the silencing construct were inoculated into wheat seedling leaves at the two-leaf stage as described previously ^78^. BSMV:PDS was used as a positive control and BSMV:γ as a negative control ^79^. At 12 days after viral infection, leaves were collected for gene expression analysis and photographing. The plants were subjected to drought stress when symptoms in BSMV were visible. Photographs were taken after 15 days of drought, and survival statistics were calculated. Primers used in the experiments are listed in Extended Data Table S15.

### Introgression of the *AetHap2* allele

A group of *Ae. tauschii*-wheat introgression (A-WI) lines were developed from the cross between *Ae. tauschii* accession T093 (2*n* = 14, DD) and the common wheat cultivar Zhoumai 18 (2*n* = 6*x* = 42, AABBDD) ^80^. KS262 is a key BC3F8 line carrying only one introgression segment on chromosome 2D, based on evaluation by 55K SNP array. To assess the genetic effect of introgression segments, a derived NIL population was developed by crossing KS262 with Zhoumai 18.

### DAB and NBT staining

DAB and NBT staining were performed following the stain kit manual. Briefly, wheat seedlings were grown in soil under normal conditions for three weeks, and water was withheld for another week before plants were sampled. Leaves were stained in 1 mg/mL DAB solution (Coolaber, SL1805) or 1 mg/mL NBT solution (Coolaber, SL1806) in the dark for 12 h at 37°C. After staining, the leaves were bleached with 85% ethanol in a water bath at 95°C until chlorophyll completely faded, and then photographed using a Nikon D80 camera.

### Transcription anti-termination assay in *E. coli*

The nucleic-acid unfolding activity of TaCSP-H1 and its mutations were measured as previously described with modifications ^81^. The cDNA of *TaCSP-H1D* was fused with pINIII vector and transformed into *E. coli* RL211 strain. The pINIII vector containing *CspA* and pINIII empty vector were used as positive and negative controls, respectively. Transformants were cultured in LB complemented with 50 μg/ml ampicillin. When the culture grew to an OD600 nm of 0.4-0.6, 1 mM IPTG was added, and cultivation continued for another 2 h. The cultures were diluted 10-fold serially and spotted on LB plates containing 50 μg/ml ampicillin and 1mM IPTG, with or without 15 μg/ml chloramphenicol. The plates were incubated at 37°C for 2-3 days and then photographed.

### Complementation assay of *E. coli* BX04

The complementation of *E. coli* BX04 strain (*ΔcspAΔcspBΔcspEΔcspG*) was performed as previously described ^35^. Briefly, the cDNAs of *TaCSP-H1D, TaCSP-V1D, TaCSP-V2D,* and *TaCSP-V3D* were respectively fused with pINIII vector and transformed into *E. coli* BX04 cells. As a positive control, the pINIII vector containing *CspA* was also constructed. *E. coli* strain BX04 cells transformed with different plasmids were grown on LB medium, and 10-fold serial dilutions of the culture were spotted onto LB plates and incubated at 37°C or 15°C.

### *In vitro* nucleic-acid unfolding assay

A fluorescent molecular beacon system was used to examine the nucleic-acid unfolding activity of TaCSP-H1 *in vitro* as described previously ^82^. The beacon used in this study was a 78-nt hairpin-shaped molecule labeled with a fluorophore and quencher: tetramethyl rhodamine-AGGGTTCTTTGTGGTGTTTTTATCTGTGCTTCCCTATGCACCGCCGACGACAGTCGCTAACCTCTCGCTAAGAACCCT-DABCYL; the molecular beacon was synthesized by Sangon Company. The fluorescence was determined by an F-4700 FL spectrofluorometer (Hitachi). The excitation and emission wavelengths used were 555 nm and 575 nm, respectively. The fluorescence of a 500-μL solution of 30 nM molecular beacon dissolved in reaction buffer (20 mM Tris-HCl, pH7.5, 1 mM MgCl_2_) was monitored after adding each protein.

### Measurement of the agronomic traits and yields

Seeds of *TaCSP-H1-*OE lines, *tacsp-h1* mutants and Fielder were treated at 4°C for three days and placed at room temperature for germination. After one week, the seeds were moved into plastic pots containing an equal amount of soil, and the pots were watered with the same amount of water to keep the soil water content at 30%. Seedlings were divided into two groups for drought and well-watered treatments. The well-watered treatment group was watered timely to keep the soil water content at 30%, while the soil water content of drought treatment group was held at 8%. Watering was terminated at the beginning of senesced yellowing. Agronomic traits and grain yields were measured after harvesting.

## Reporting summary

Further information on research design is available in the Nature Portfolio Reporting Summary linked to this article.

## Data availability

The RNA-seq and RIP-seq data generated in this study have been deposited in the National Center for Biotechnology Information (NCBI) under BioProject number PRJNA1114877. All analyses were conducted using standard software. The settings of software used for analyses were described in the Methods. Materials used in this study are available upon request. Source data are provided with this paper.

## Acknowledgments

We thank Suoping Li for providing the T093-Zhoumai18 introgression lines. We also thank Wei Chi for providing the *E. coli* BX04 strain and PINIII vector, and Xiuzhu Dong and Jie Li for providing the *E. coli* RL211 strain. This work was supported in part by grants from the National Natural Science Foundation of China (32230079), National Key Research and Development Program of China (2022YFF1001602), Key Research and Development Program of Henan Province (231111110200), and Key Research Project of the Shennong Laboratory (SN01-2022-01).

## Author contributions

C.-P.S., Y.Z. and J.H. conceived and designed the study. K.W., G.G., S.B., J.M. and Z.Z. performed main experiments. J.H., J.M. and S.B. performed HGT analyses. K.W., G.G., Z.Z., Z.X., J.W., J.L. and X.Z. generated the transgenic lines and identified the phenotypes. Z.X., J. W., W. X. and C.Y. performed the field tests and population genetic analysis. W.W. analyzed the protein structures. Z.X., W. X., Z.Z., H.L., Z.L. and Y.L. collected the introgression lines data. C.-P.S., Y.Z., J.H., K.W. and S.B. analyzed the RNA-seq and RIP-seq data. T.B., W.-C.L., H.L., X.S., R.-F.S., J.L., Q.L., F.Z., X.Q. and T.H. conducted some of the experiments. C.-P.S., Y.Z., J.H. and K.W. wrote the manuscript. C.-P.S., Y.Z., J.H., K.W., G.G. and S.B. revised the manuscript.

## Competing interests

The authors declare no competing interests.

## Notes

### Competing Interest Statement

The authors have declared no competing interest.

